# An attempt at restoring original radiocarbon ages in collagen from bioapatite

**DOI:** 10.1101/2022.11.14.516521

**Authors:** Mário André Trindade Dantas, Alexander Cherkinsky

## Abstract

In the literature there is a consensus that radiocarbon dating performed in bioapatite presents younger datings than those performed in collagen, thus, we propose a general regression that could be used to convert the radiocarbon dating performed in bioapatite to the original ones in collagen in fossil samples all of the world. This general regression presents several good indexes of quality, high correlation (R^2^ = 0.98), lower values of percent predicted error (%PE = 0.01), and the standard error of the estimate (%SEE = 25), showing that is a good tool, as the predicted values are similar to those observed. Using this regression we converted the radiocarbon datings in bioapatite to collagen made for several taxa from the Brazilian Intertropical Region, and suggest that these datings could be 1-7 Cal BP kyr older than previously thought.

## 1. Introduction

For late Quaternary researchers, known the age of their samples is very important to help them in interpretations about, for example, the paleoecology and extinction of the studied taxa (e.g. Barnosky & Lindsay, 2010; Dantas et al., 2020).

These researchers have as options for direct dating the use of radiocarbon dating (AMS) in collagen (^14^C_collagen_), which could date samples until ∼60 kyr (Cook & Plicht, 2007). However, in Tropical regions they face one main problem, the lost of collagen due the diagenetic process (Hedges, 2002).

In the absence of collagen Cherkinsky (2009) presented an option in the absence of collagen, to perform the radiocarbon dating in bioapatite (^14^C_bioapatite_), justifying that the mineral fraction survives much better than organic ones, suffering small changes through the diagenesis.

Since then, several papers dealing about the chronology and paleoecology of the meso-megamammals from the Brazilian Intertropical Region has been made using this technique, and presenting the occurrence of this fauna in the Late Pleistocene, between 9-32 Cal BP Kyr (e.g. Dantas et al., 2017; 2020; 2022).

However, Some authors (Zazzo & Saliège, 2011; Zazzo, 2014) suggests that during diagenesis bioapatite exchange carbon with a ^14^C-enriched (i.e. younger) carbon source, which promote in ^14^C_bioapatite_ younger datings than ^14^C_collagen_, as older the dating, major was the difference between them. Thus, Zazzo (2014) recommended that ^14^C_bioapatite_ should be considered as minimum estimates.

Based on this observation, we propose, and test, regressions that could convert the radiocarbon dating in bioapatite to collagen in samples collected in different climatic zones (boreal, temperate, subtropical, and tropical).

## 2. Material and methods

In this paper we use several radiocarbon dating performed in collagen and bioapatite in the same samples, the collagen radiocarbon datings had expected C/N pattern (∼3) and more than 5% of collagen in each samples. Another index was the presence of modern carbon (pMC) in the samples, were used samples with proportional lower pMC as older the samples (Cherkinsky, 2009; Zazzo, 2014 and references therein; Cherkinsky et al, 2015; Table S1).

Reduced major axis (RMA, Model II) regressions were produced using the entire sample to create a general regression, and specific ones for each climatic zone (boreal, temperate, subtropical, and tropical; Table 1), because: (i) it deals better with extrapolation than ordinary least squares; (ii) incorporate an assumption that there is an error in *X*; and (iii) is symmetric, meaning that the slope of the line do not differs depending upon which variable is identified as *X* and which is *Y* (OLS, Model I; Smith, 2009; Halenar, 2011 and references therein). This method use the slope (*b*_OLS_) find in OLS, the mean values of *x* and *y*, and the absolute value of the correlation of pearson (*r*) to estimate a new slope (*b*_RMA_; equation 1) and intercept (*a*_RMA_; equation 2) (Harper, 2016).

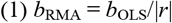

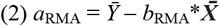

**Table 1.** Values of the RMA regressions, coeficient of determination (R^2^), average percent prediction error (%PE) and standard error of the estimate (%SEE) obtained for each climatic zone (cz).

| cz | Samples | Slope | Intercept | $R^2$ | %PE | %SEE |
| --- | --- | --- | --- | --- | --- | --- |
| Boreal | 8 | 1.10 | -0.41 | 0.97 | 0.02 | 10.90 |
| Temperate | 7 | 1.15 | -0.50 | 0.98 | 0.05 | 89.32 |
| Subtropical | 8 | 0.99 | 0.05 | 0.99 | 0.06 | 21.26 |
| Tropical | 5 | 0.97 | 0.13 | 0.99 | 0.00 | 25.16 |
| All | 28 | 1.09 | -0.31 | 0.98 | 0.01 | 25.00 |

As the radiocarbon datings do not presented a normal distribution (Shapiro-Wilk test, *p* < 0.05), these data were transformed to logarithm values (at base 10) to approximate a log-normal distribution, due it assigns equal weight to all data points in a regression (e.g. Smith, 1993 and references therein).

In addition to the correlation of data log-transformed, as high correlation do not means that the regression is a good predictor (e.g. Smith, 1984), were calculated the percent predicted error (%PE) and the standard error of the estimate (%SEE).

The %PE of each sample is calculated using equation (3) (Van Valkenburgh, 1990 and references therein; Halenar, 2011), and then is made an average of the absolute %PE mean of the variables. This index provides a comparative value to see the predictive accuracy of the regressions.

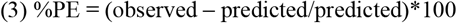

To estimate the %SEE we use the equation (4), this index reflects the ability of the independent variable to predict the dependent variable (Van Valkenburgh, 1990 and references therein). SE is the standard error (= standard deviation/√n).

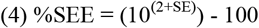

To test if are statistical differences between the proposed regressions was made by the analysis of variance, we used ANOVA (1 factor, α = 0.05) in PAST 3.11 software (Hammer et al., 2001).

The best estimated regression (results and discussion) was used to correct the radiocarbon dating in bioapatite to collagen of eight extinct meso-megamammals from the Brazilian Intertropical Region (BIR; sensu Cartelle et al., 1999; Table 2).

**Table 2.**
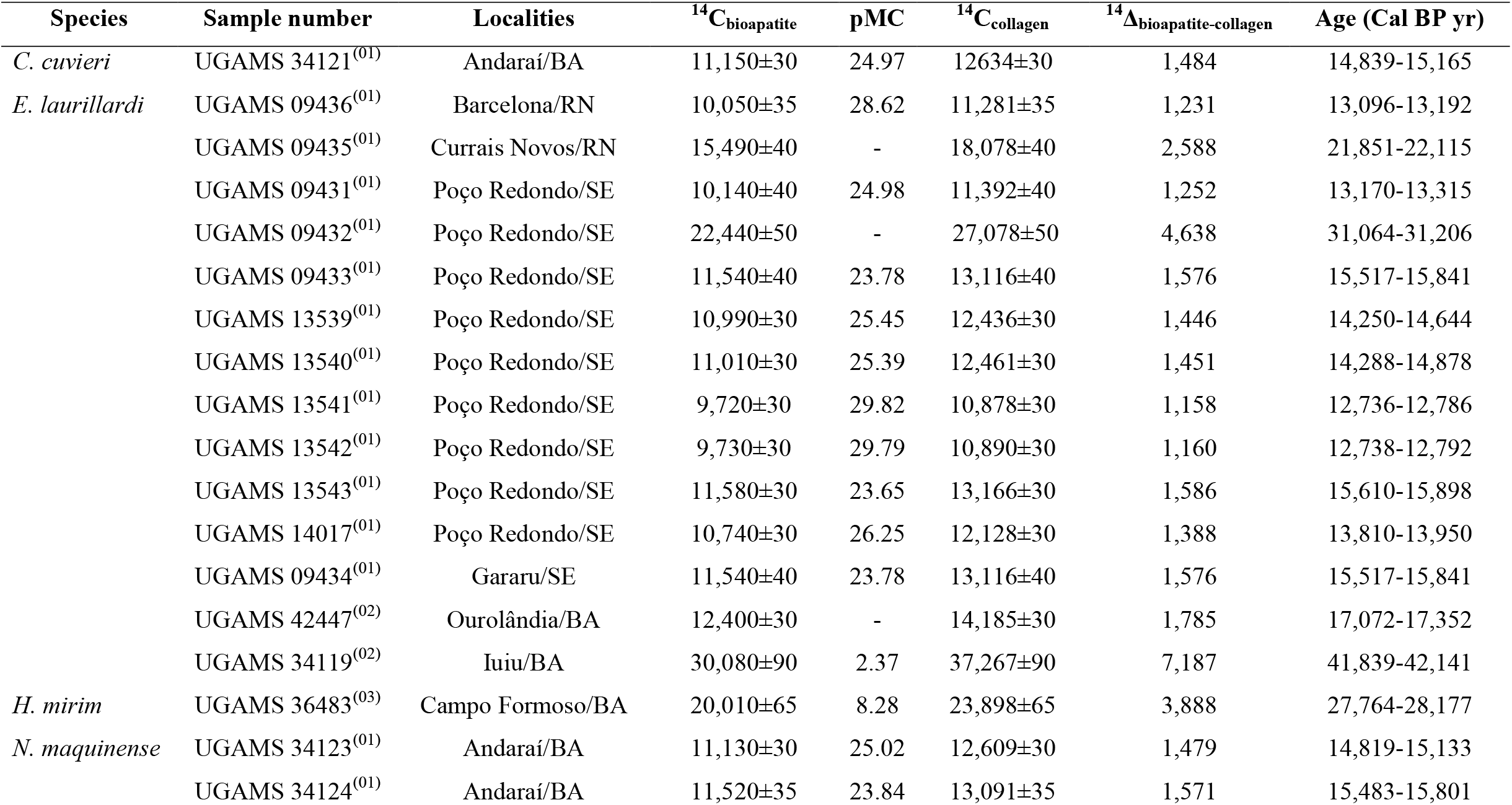
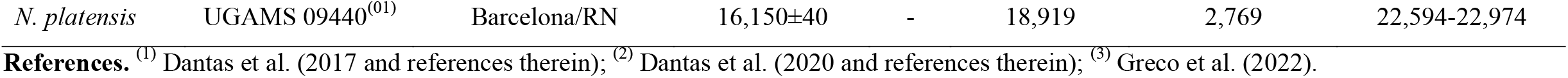
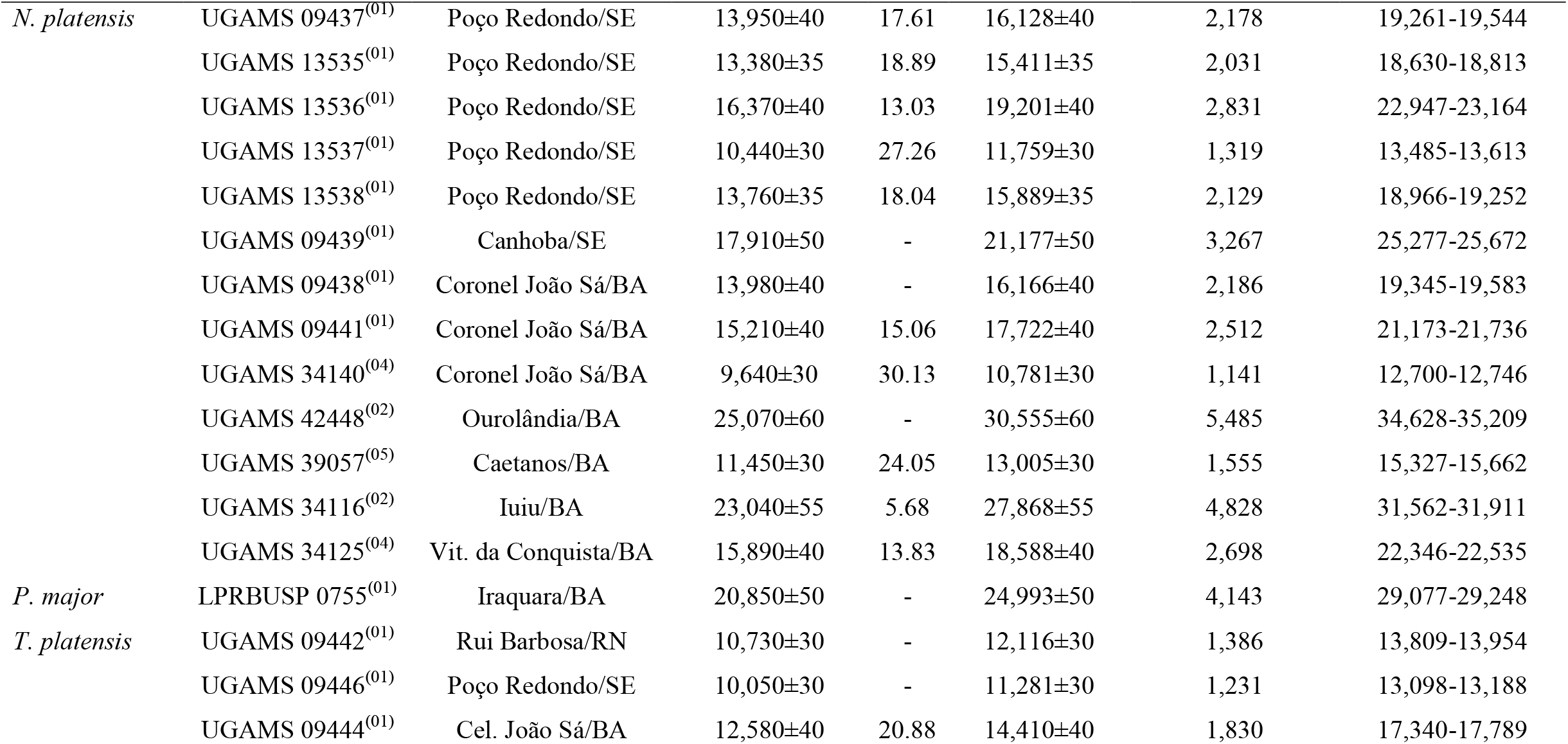
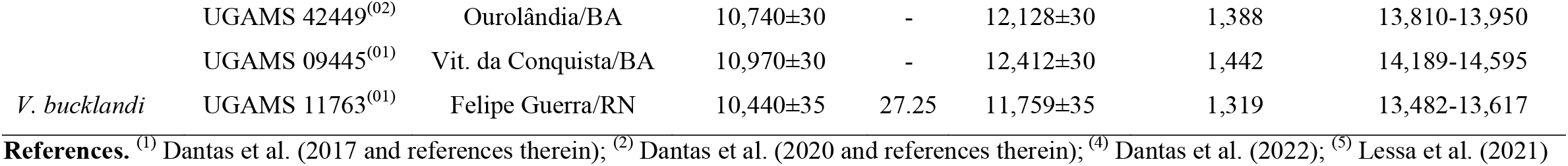
Radiocarbon datings in bioapatite (^14^C_bioapatite_) converted to collagen (^14^C_collagen_), presence of modern carbon (pMC), and calibrated ages (SHCal20 curve) for extinct Late Pleistocene meso- megamammals taxa from Brazilian Intertropical Region.

## 3. Results and discussion

### 3.1. Converting ^14^C_bioapatite_ into ^14^C_collagen_

The radiocarbon dating samples (dated both in bioapatite and collagen) came from different countries located in boreal, temperate, subtropical, and tropical climatic zones (Table S1), in all those were observed that the diagenesis altered the bioapatite and provide, in general, younger radiocarbon dating in comparison with those found in collagen (Cherkinsky, 2009; Zazzo, 2014 and references therein; Cherkinsky et al, 2015).

Using these data were estimated regressions for each climatic zone, plus, a general one, and noted that they are similar (ANOVA, *F*_*obs*_ = 0.02, *p* = 0.99; Table 1), presenting strong correlations and similar slopes (m) values, however, presents different %PE and %SEE.

The slopes of these RMA regressions, created with the available data, allow us to interpret that the radiocarbon dating in bioapatite tend to be slightly lower than that in collagen in Boreal (m = 1.10), Temperate climate zones (m = 1.15), and in all world (m = 1.09). In Subtropical climate zones (m = 0.99) and Tropical climate zones the radiocarbon dating in bioapatite tend to be slightly higher than that in collagen (m = 0.97).

If was choose to use the regressions for each climatic zone there is a tendency to find different corrected collagen datings, being that for Temperate climatic zones higher than in the others zones. To avoid this, as all regressions are similar, the general regression must be used (Figure 1), as it presents a strong correlation (R^2^ = 0.98), lower mean %PE (= 0.01; Table 1), and average %SEE (= 25.00; Table 1), combinated.

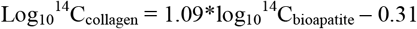

**Figure 1.**
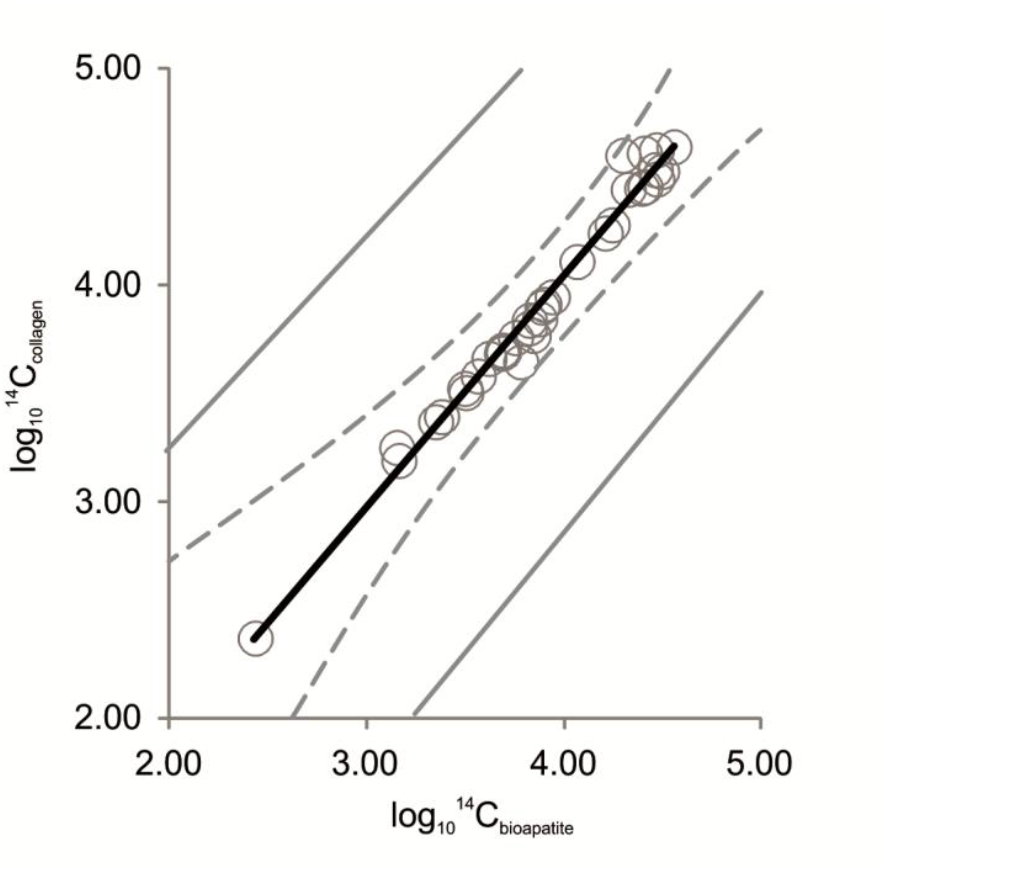
Reduced major axis Regression of log Radiocarbon dating (bioapatite) and log Radiocarbon dating (collagen) using 28 samples (Table S1). Regression line (Black solid line), confidence intervals (Gray dotted lines), and prediction intervals (Gray solid lines).

The best regressions must have higher values of correlation, lower values of %PE (<15%) and %SEE (Delson, et al., 2000; Ruff, 2003), showing that the predicted values are similar to those observed, which this general equation reached.

### 3.2. Limit of convertion

The radiocarbon dating calibration curve could allow estimating the age of terrestrial samples to about 50 kyr, the limit of the method (Cook & Plicht, 2007; Wood, 2015). This limit depends on the pretreatment used, which could help to better purify the samples from contaminants, and allows older datings (Wood, 2015).

As stated before the radiocarbon dating in bioapatite is considered as minimum ages, and the proposed general regression can convert the ^14^C_bioapatite_ to ^14^C_collagen_, however observing the limit of the method (50 kyr) this regression should be used to convert only ^14^C_bioapatite_ ∼39,400 yr. Older converted collagen dating could be not calibrated in CALIB 8.1 program (Reimer et al., 2020) due to the extrapolation of the limit of 50 kyr.

### 3.3. Study case: converting the ^14^C_bioapatite_ of the meso-megamammals from the Brazilian Intertropical Region

Using the new general regression, were converted the radiocarbon datings made in bioapatite to the collagen for eight extinct meso-megamammals taxa which lived in the BIR, and later calibrated into calendar ages before present, using the same standard error found in the ^14^C_bioapatite_, using CALIB 8.1 program (Reimer et al., 2020), SHCal20 curve (Hogg et al., 2020), and 2σ measured ages reported in Table 2.

The difference between the radiocarbon dating in bioapatite to the converted to collagen shows a variation between 1,141 to 7,187 years (Table 1), while the difference between the calibrated datings was 1,166 to 7,523 Cal BP yr older than previously thought (e.g. Cherkinsky et al., 2013; Dantas et al., 2017; Figure 2).

**Figure 2.**
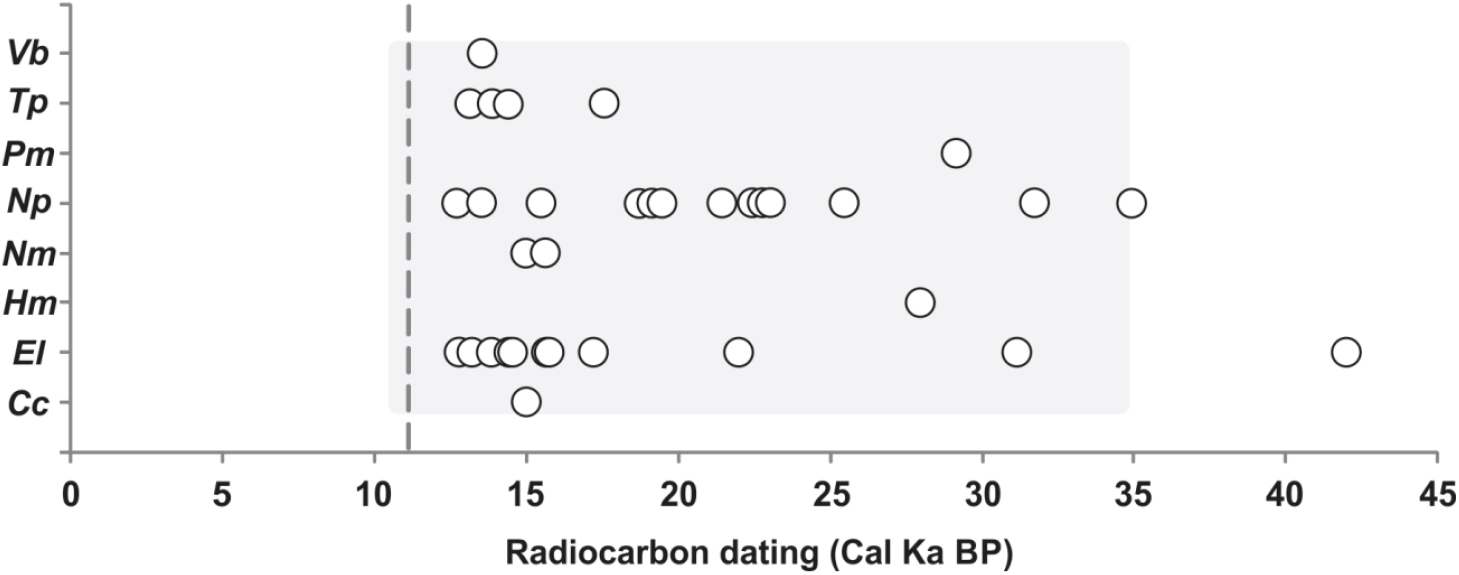
Radiocarbon Chronology in collagen (white circles) of extinct meso-megamammals from the Brazilian intertropical region. Gray shadow represents the interval found in Radiocarbon dating performed in bioapatite. Doted Gray line represents the limit between Pleistocene-Holocene. **labels:** *Cc* - *Catonyx cuvieri*; *El* - *Eremotherium laurillardi*; *Hm* – *Hemiauchenia mirim*; *Nm* – *Nothrotherium maquinense*; *Np* - *Notiomastodon platensis*; *Pm* - *Palaeolama major*; *Tp* - *Toxodon platensis*; *Vb* – *Valgipes bucklandi*.

The diagenesis could promote small alterations in ^14^C/^12^C in bioapatite carbonate, leading to younger datings, however, this alteration is non-significant in ratio stable isotopes of carbon (^13^C/^12^C), at least, for the last 40 thousand years (Zazzo, 2014).

When the diagenesis affect the bioapatite, the substitutions are mainly in the hydroxyl position in the phosphate, and even with a carbonate substitution occurs, the isotope signature in stable and radioactive carbon maintain the original signature (Cherkinsky, 2009).

The available *δ*^13^C associated to the converted ^14^C_collagen_ for the megafauna of the BIR brings paleoecological information of a time span ranging ∼12,700 to 42,100 years (Figure 2), and allow suggesting that these meso-megamammals lived in the BIR, at least, until 12 kyr, in the Late Pleistocene. Considering other dating techniques, as for example Electron Spin Ressonance, this time span could be expanded to 9±2 ky (Ribeiro et al., 2013).

## 4. Final remarks

In this paper was proposed a regression to convert the radiocarbon dating performed in bioapatite to collagen, allowing facilitating the comparison of radiocarbon datings in all world.

Using this new tool were converted the radiocarbon dating performed in bioapatite in fossils of meso-megamammals from Brazil and suggest that these datings are 1-7 Cal BP kyr older than previously thought.

## Acknowledgements

To CNPq for the research fellowship for MATD [PQ/CNPq 311003/2019-2]. To MSc. Lais Alves Silva for the critical review of the manuscript.

